# Magnetogenetic stimulation inside MRI induces spontaneous and evoked changes in neural circuits activity in rats

**DOI:** 10.1101/2023.12.14.571681

**Authors:** Kai-Hsiang Chuang, Chunqi Qian, Assaf Gilad, Galit Pelled

## Abstract

The ability to modulate specific neural circuits and simultaneously visualize and measure brain activity with MRI would greatly impact understanding brain function in health and disease. The combination of neurostimulation methods and MRI in animal models have already shown promise in elucidating fundamental mechanisms associated with brain activity. We developed an innovative magnetogenetics neurostimulation technology that can trigger neural activity through magnetic fields. Similar to other genetic-based neuromodulation methods, magnetogenetics offers cell-, area- and temporal-specific control of neural activity. However, the magnetogenetics protein (Electromagnetic Preceptive Gene (EPG)) are activated by non-invasive magnetic fields, providing a unique way to target neural circuits by the MRI gradients while simultaneously measure their effect on brain activity. EPG was expressed in rat’s visual cortex and the amplitude of low-frequency fluctuation (fALFF), resting-state functional connectivity (FC), and sensory activation was measured using a 7T MRI. The results demonstrate that EPG-expressing rats had significantly higher signal fluctuations in the visual areas and stronger FC in sensory areas consistent with known anatomical visuosensory and visuomotor connections. This new technology complements the existing neurostimulation toolbox and provides a mean to study brain function in a minimally-invasive way which was not possible previously.

## Introduction

The neural circuit of the brain underpins our behavior and cognition. The capability of modulating a targeted circuit lies great potential for improving cognitive performance or relieving disease symptoms. The expansion of molecular and biological tools to modulate neural function can lead to improved and novel approaches to deliver neuromodulation that can offer cell-precise, temporally precise, region-specific, and non-invasive way to manipulate cellular function. The ability to modulate specific neural circuits and simultaneously visualize and measure brain activity would greatly impact understanding brain function in health and disease. MRI is a powerful tool for mapping brain-wide neural activity and connectivity induced by targeted neural activation. Developing technologies that will allow to trigger or suppress neural activity of a specific population of cells, while simultaneously acquiring neural activity via MRI methods remains an active area of research. Over the past decade, the combination of MRI with other neurostimulation methods such as implantable electrodes [1], optogenetics [2-6], chemogenetics [7] and transcranial magnetic stimulation [8] have elucidated several fundamental neural mechanisms including hemodynamic signals, plasticity mechanisms, resting-state fMRI, memory consolidation, and epilepsy.

A new neuromodulation method that can trigger neural activity through magnetic fields would provide an innovative and accessible way to manipulate neural circuits *in vivo*. Magnetogenetic*s* is the field of manipulating cells by magnetoreception-inspired bioengineering of novel genetic tools. For example, radio-waves and magnetic fields used for heating iron nanoparticles conjugated to ion channels were used to remote control cellular activity [9, 10]. Other studies reported on engineering two artificial chimeric magneto-sensors where ion channels were fused to ferritin [11, 12]. Nevertheless, the ability to sense and respond to magnetic fields is well documented in many species, especially in diverse groups of fishes[13]. Recently, we discovered a novel *electromagnetic perceptive gene* (EPG) from the glass catfish *Kryptopterus vitreolus* [14, 15]. Recent work demonstrated that EPG, a 13.3 kD protein, is anchored via glycosylphosphatidylinositol (GPI) anchor to the cell membrane facing the extracellular space and increased intracellular calcium upon activation with electromagnetic field (EMF) [16]. EPG may undergo conformational changes when exposed to an EMF [17]. Calcium imaging in cell culture [18] and electrophysiology recordings in rat brain slices [19] have demonstrated that EMF stimulation of EPG-expressing neurons leads to a significant response. In these experiments, an electromagnet device delivering 50-150 milli-Tesla (mT) was positioned over the cell culture and the brain slice to deliver the magnetic stimulation [20]. Furthermore, it was demonstrated that this technology can induce behavioral changes at a level comparable to more conventional neurostimulation methods. For example, in a rat model of peripheral nerve injury, rats that expressed EPG in excitatory neurons in the somatosensory cortex showed improved sensorimotor performance when they were subjected to a magnetic stimulation [21]; In a rat model of temporal lobe epilepsy, rats that expressed EPG in inhibitory interneurons in the hippocampus showed less seizures when injected with kainic acid[19]. Thus, magnetogenetics could provide cell-specific, area-specific and temporal-specific features that a neuromodulation strategy requires, with the advantage of minimal invasiveness and technical challenges associated with electrophysiology or illumination inside an MRI scanner.

The EPG has been shown to be active in response to local magnetic fields in the order of mT. Nevertheless, it remains unclear how EPG activation may affect the activity and functional connectivity of specific brain circuits, and what is the range of magnetic fields that activates the EPG. The former is essential for understanding how EPG activation alters brain function and associated behavior. The latter would be crucial to continue and artificially design and synthesize new EPGs that could be sensitive and tunable to different magnetic field strengths and thus useful for different biological and physiological applications. In this study, we evaluated how EPG expression in the rodent’s visual cortex would influence the amplitude of low-frequency fluctuation (fALFF), resting-state functional connectivity (FC), and sensory activation using a 7T MRI. We found that EPG activation by MRI increased fluctuation amplitude and connectivity between cortical and subcortical areas. Furthermore, visual evoked responses in EPG-expressing rats were larger compared to controls. Thus, magnetogenetics will allow inducing or silencing neural activity within an MRI. It would open a new array of possibilities to study brain circuits in health and disease.

## Methods

### Animal preparation

All procedures were approved by the Institutional Animal Care and Use Committee at Michigan State University. Adult male and female Long Evans rats (200-400g, n=10, 6 females, 4 males) rats were anesthetized with isoflurane (5% for induction; 2.5% for maintenance and surgery) and positioned in a stereotaxic frame. Stereotaxic injections of 5µl at each location contained AAV1-CaMKIIα - {EPG(Rat)X3Flag}:IRES:EGFP at a titer of 10^13^ GC/ml (n=5). Sham-control rats were injected with AAV-CaMKIIα-EGFP (n=5). See supporting information for DNA sequence detail. The microinjector was positioned at 4 location in the visual cortex: AP -7.3mm, ML 3.2mm; AP -7.3mm, ML -3.2mm; AP -5.4mm, ML 4mm; AP -5.4mm, ML -4mm. Two to four weeks after stereotaxic injections MRI measurements took place. The EPG sequence can be found in the gene bank: *(GenBank: MH590650*.*1)*. For MRI the anesthesia was induced with 5% isoflurane, followed by a bolus injection of dexmedetomidine (0.05 mg/kg, sc; Dexdomitor®, Orion Pharma), after which isoflurane was discontinued, and a constant dexmedetomidine infusion (0.1 mg/kg/hr) was administered subcutaneously. During fMRI measurements, rats’ temperature was maintained at 37°C, and the breathing rate, partial oxygen saturation, and heart rate were continuously monitored.

### MRI data acquisition

Data acquisition was performed using a 7T MRI system (Bruker BioSpec, Germany) with a volume coil for transmission and a 4-channel brain array (Bruker T11483V3) for reception. High-resolution anatomical scans were acquired using T2-weighted fast spin echo with a spatial resolution of 100x100x650 µm and with TR=4.2 s, TE=24 ms, RARE factor=4, Slice Number=32. Functional scans were acquired using single-shot gradient-echo echo planar imaging with TR=1000 ms, TE= 20 ms 40x40 matrix and 32 slices with a spatial resolution of 650x650x650 µm. Resting state fMRI scans were acquired over 12 minutes (720 scans). Afterwards, two kinds of sensory stimulations were used. The first was forepaw stimulation for verifying that the functional response is in the appropriate location, such as primary somatosensory cortex forelimb area (S1FL). The second was visual stimulation to evaluate the influence of the visual EPG on the visual activation. The electrical forepaw stimulation consisted of a 3 Hz pulse train of 0.5 mA applied to the right forepaw. Visual stimulation was delivered to both eyes using fiber optic cables presenting 5 Hz flashing lights. Both kinds of stimulations had alternating ON/OFF block design starting with 30-second OFF, then alternating 20-second ON and 20-second OFF for two times, leading to 110 total scans.

### MRI analysis

The MRI data were processed using MATLAB (MathWorks Inc), FSL (v6.0, https://www.fmrib.ox.ac.uk/fsl), AFNI (ver 18, National Institutes of Health, USA) and ANTs (v2.3.1, http://stnava.github.io/ANTs). The fMRI preprocessing followed an optimized pipeline we developed [22]. The first 5 scans were discarded to ensure that the baseline signal reached steady state. Potential motion artifact was corrected by FSL mcflirt, followed by brain extraction automatically using PCNN3D [23] and manual editing. Nuisance signals, including quadratic drift, 6 motion parameters and their derivatives, 10 principal components from tissues outside the brain which included muscle and scalp were extracted [22]. The time-series intensity of each voxel was normalized by the mean signal of the first OFF period (30 s) to convert the BOLD signal into percent signal change. The data was coregistered to the 0.3-mm SIGMA rat brain template [24] via the structural T2-weighted MRI by linear and nonlinear transformations using ANTs. The data were then smoothed by a 1.0 mm 3D Gaussian kernel. The stimulus-evoked fMRI data were high-pass filtered at 0.01 Hz to account for any potential baseline fluctuation. A general linear model was used to detect the evoked activation by convoluting the stimulus paradigm with a rodent hemodynamic response function [25] and included the nuisance signals as confound regressors in the design matrix. The fitted coefficient beta was regarded as the activation level in voxel-wise and ROI-wise statistics.

For resting-state functional connectivity, the nuisance signals were first regressed out and the data were band-pass filtered at 0.01-0.1 Hz. To avoid the filter artifact, the first and last 15 time points were removed. Seed-based correlation analysis was used to measure FC across the brain. Based on the SIGMA template, the brain was divided into 170 bilateral gray matter regions-of-interest (ROIs) over the whole brain (see supplementary Table S1 for a complete list of brain regions and abbreviations). The averaged time-course of each brain region was extracted as a seed signal. Pearson’s correlation coefficients between time-courses in each voxel and ROI were calculated using AFNI 3dNetCorr. Fisher’s *z*-transformation was used to convert correlation coefficients to *z* values. Furthermore, the amplitude of low frequency fluctuation (ALFF) was used to measure the spontaneous activity change induced by EPG. The spectral power of voxel time-course with nuisance removal but without band-pass filter was calculated. The total power within the 0.015 to 0.2 Hz range was used as the ALFF. The fractional ALFF (fALFF) was calculated by normalizing the ALFF by the total power above 0.015 Hz.

### Immunohistochemistry

Rats were perfused transcardially with 1X Phosphate Buffered Saline (PBS) and 4% Paraformaldehyde (PFA). The brains were extracted and placed in 4% PFA overnight after which they were placed in varying concentrations of sucrose. Frozen brains were sectioned at a thickness of 50µm and slices were placed in 4C in PBS. Slices were washed in PBST and a blocking solution added (donkey serum, Sigma-Aldrich, St. Louis, MO). Slices were then incubated overnight with the primary antibodies (1:500, Anti-CamKII, ab52476, and Anti-GFP, ab13970, Abcam, Boston, MA). The following day, slices were washed and incubated with the secondary antibodies (1:500, Donkey Anti-Rabbit conjugated with Alexa Fluor 657, and Donkey Anti-Chicken conjugated with Alexa Fluor 488, Jackson ImmunoResearch Labs, West Grove, PA). Images were acquired using the Biotek Cytation 5 Cell Imaging Multi-Mode Reader (Biotek Instruments, Inc., Winooski, VT) configured on an inverted microscope.

### Statistical analysis

Within and between group comparison was conducted in each voxel or ROI. Voxel-wise group comparison was calculated by one-sample t-test and thresholded at p<0.01 (False Discovery Rate [FDR] corrected). Between-group differences were calculated by two-sample t-test and thresholded at p<0.05 (FDR corrected). For the evoked fMRI, the beta maps were compared, and for the resting-state FC the z-score maps were compared. The resting-state fALFF of each ROI was compared using 2-way ANOVA (Prism 9, GraphPad Software). For visualization, significant connections were overlaid on the 3D-rendered brain atlas using BrainNet Viewer (https://www.nitrc.org/projects/bnv/).

## Results

Resting-state fMRI images were collected immediately after positioning the rat in the scanner to determine whether the magnetic field of MRI would activate the EPG and alter the spontaneous neural activity. The signal fluctuation amplitude and spectral power at the resting state were measured for 12 min. Unlike a typical resting-state fMRI study that only inspects fluctuations below 0.1 Hz, we observed larger changes between 0.1 to 0.2 Hz (Fig. 1A). Based on the spectral analysis, we calculated the fALFF within the 0.015 to 0.2 Hz range. Voxel-wise comparisons show that there are sparse areas showing significantly higher signal fluctuation amplitudes such as the retrosplenial cortex (RSC) (Fig. 1B). Regional analysis shows a significantly larger fALFF in the EPG group (Two-way ANOVA, F (1, 64) = 12.88, p = 0.0006) with the most significant change found in the V1m (p = 0.016, Fisher’s LSD test) (Fig. 1C).

**Figure 1.**
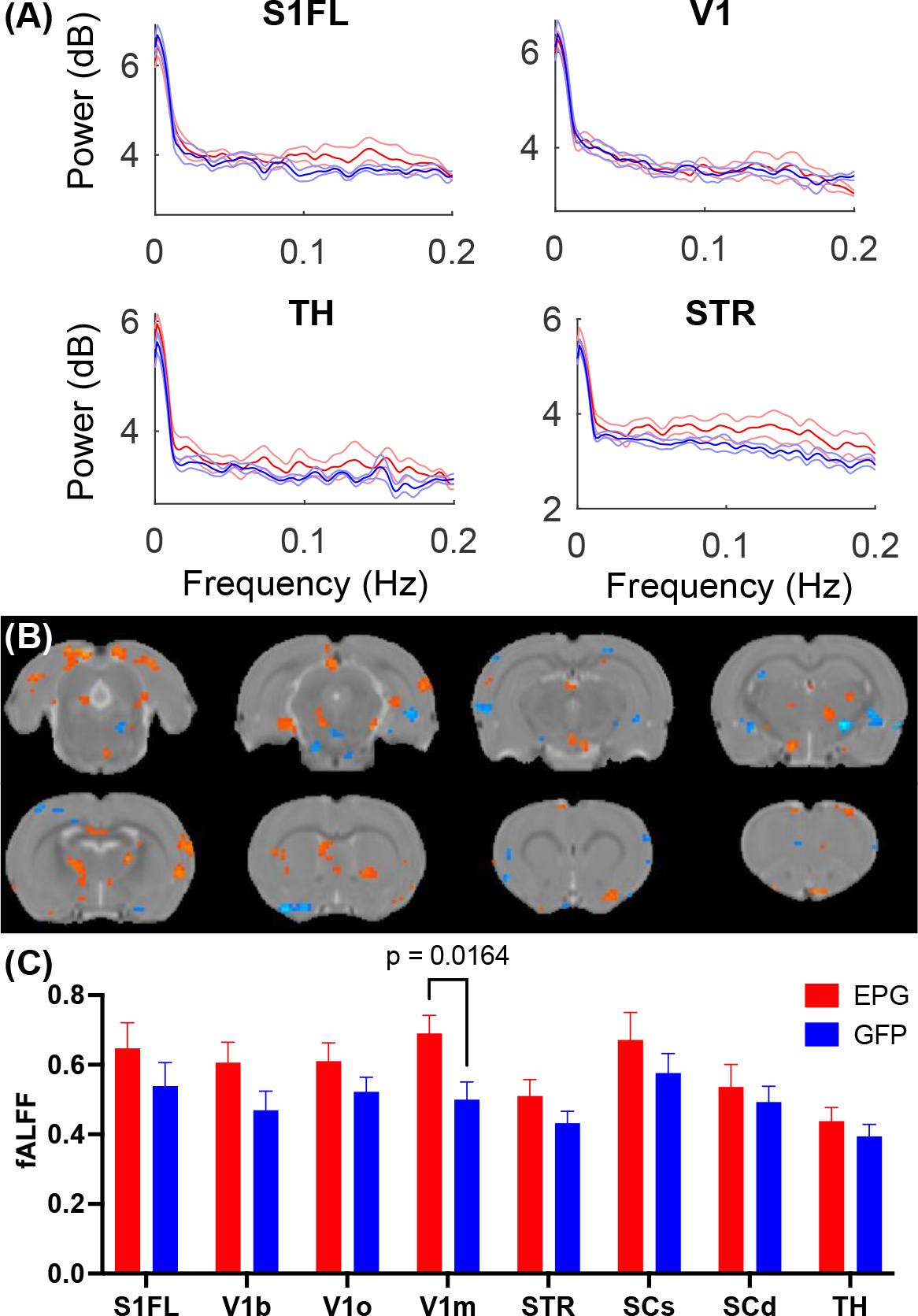
EPG expression in the visual cortex leads to increases in fractional amplitude of low frequency fluctuation (fALFF). (A) Power spectra of the resting-state fMRI signals from selected brain regions show elevated signal fluctuations above 0.1Hz in the EPG (red) compared to the GFP (blue) group. Errorbar represents the standard error of mean (SEM). (B) Voxel-wise comparison of fALFF between the EPG and GFP groups (two-sample t-test, p < 0.05, FDR corrected). (C) Regional analysis of the fALFF demonstrated that EPG rats exhibit significantly larger fluctuations at the primary visual cortex (V1m) compared to control rats (Two-way ANOVA, F (1, 64) = 12.88, p = 0.0006). This increased spontaneous neural activity in the region that the EPG was expressed suggests that EPG could be activated by the magnetic field of the MRI. See supplementary Table S1 for abbreviations of brain regions.

To further understand the effects of EPG activation on distant brain regions, we calculated the inter-regional signal correlation as a measure of FC in EPG-expressing rats. With a seed ROI at the V1m, broadly distributed FC can be seen throughout the cortex and subcortical areas, such as the RSC, S1, SC, TH, hypothalamus (Hy), and septum (Sep) (Fig. 2A, p < 0.05 FDR corrected). Whole brain analysis revealed increased FC between the cortical areas, including the primary and secondary visual, retrosplenial, entorhinal, perirhinal, primary somatosensory, auditory, motor, and orbitofrontal cortices, as well as with some subcortical areas, such as the basal forebrain (BF), SC and IC, and the cerebellum (Fig. 2B, p < 0.001 uncorrected).

**Figure 2.**
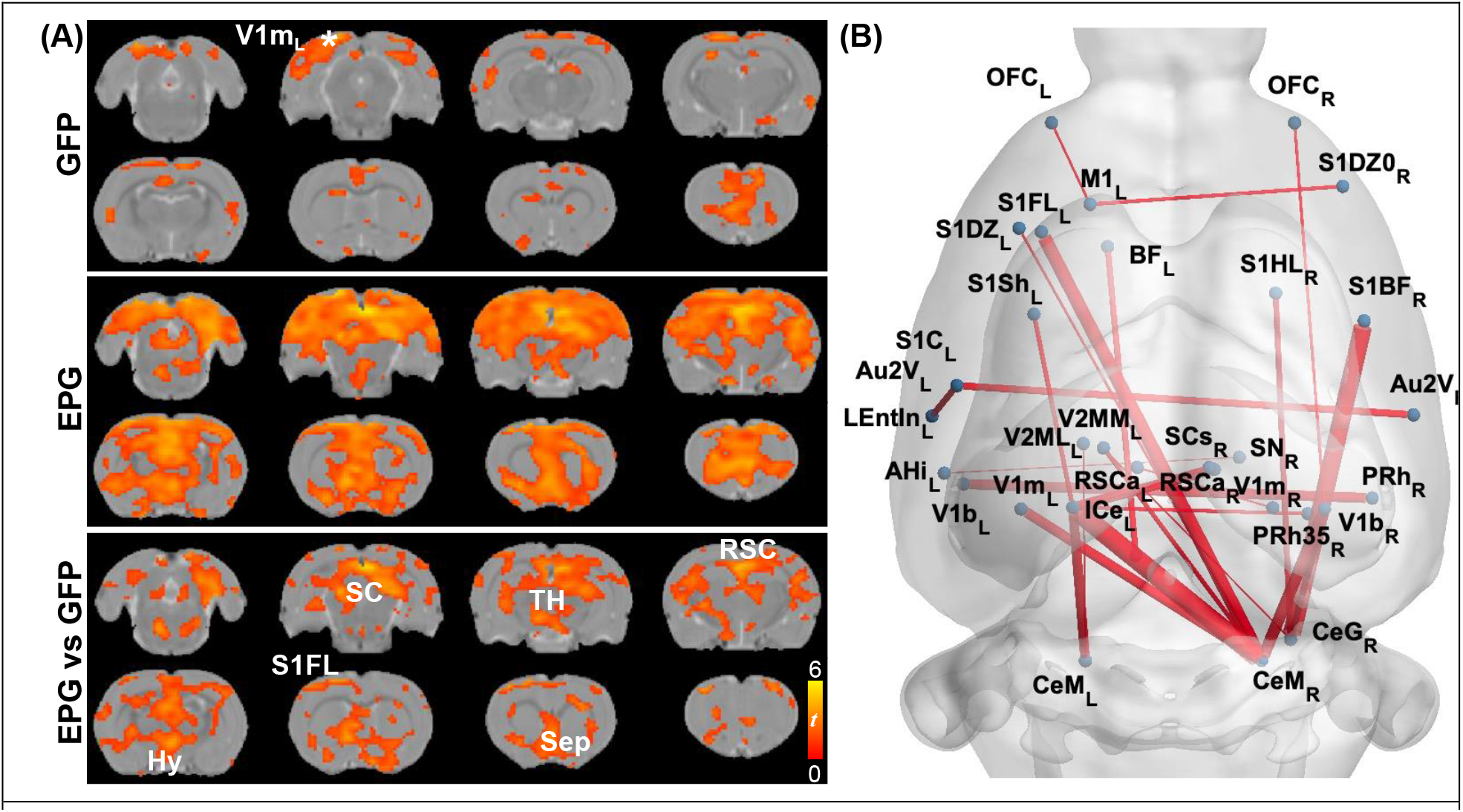
EPG expression in the visual cortex leads to increases in Functional Connectivity (FC) across cortical and subcortical areas. (A) A seed ROI was placed in the V1m of the left hemisphere, where the EPG was expressed. In the GFP control group, FC was mostly in the visual cortex with certain connectivity with the medial prefrontal cortex (one-sample t-test, p<0.01 FDR corrected). In the EPG group, broadly increased FC can be seen over the whole brain. Between group comparison shows significantly higher connectivity between brain areas of EPG rats compared to controls (two-sample t-test, p < 0.05 FDR corrected). (B) Whole brain FC (two-sample t-test, p<0.001 uncorrected) shows extensively increased connectivity beyond the visual cortex. See supplementary Table S1 for abbreviations of brain regions.

Under visual stimulation, rats expressing the control virus (GFP group; Fig. 3B) exhibit focal activation (p < 0.01, FDR corrected) in the visual pathway, including the dorsal lateral geniculate nucleus of the thalamus (LGd), superior colliculus superficial/deep subregions (SCs/SCd) to the primary visual cortex (V1) and its binocular (V1b) and monocular (V1m) subregions. Rats in the EPG group (Fig. 3C) show not only stronger activation in the visual pathway but also more distributed activation in the entire thalamus (TH). Comparing the EPG with the control group (p < 0.05, FDR corrected; Fig. 3D), stronger activation was seen in the V1, hippocampus (HP) and ventral thalamus. A decreased signal was seen in the striatum (STR), inferior colliculus (IC) and cerebellum. Interestingly, reduced activation is also seen in the primary somatosensory cortex (S1). Comparing the averaged BOLD signal time-courses in the V1, SC and TH, the responses were comparable between the EPG and GFP-control groups (Fig. 3E), indicating that the EPG did not alter the hemodynamic response function. Unlike the subcortical areas, the BOLD activation in the V1 shows a faster decrease, consistent with previous studies [26]. A slightly higher and less variable activation with a much smaller standard deviation among individuals was seen in EPG group.

**Figure 3.**
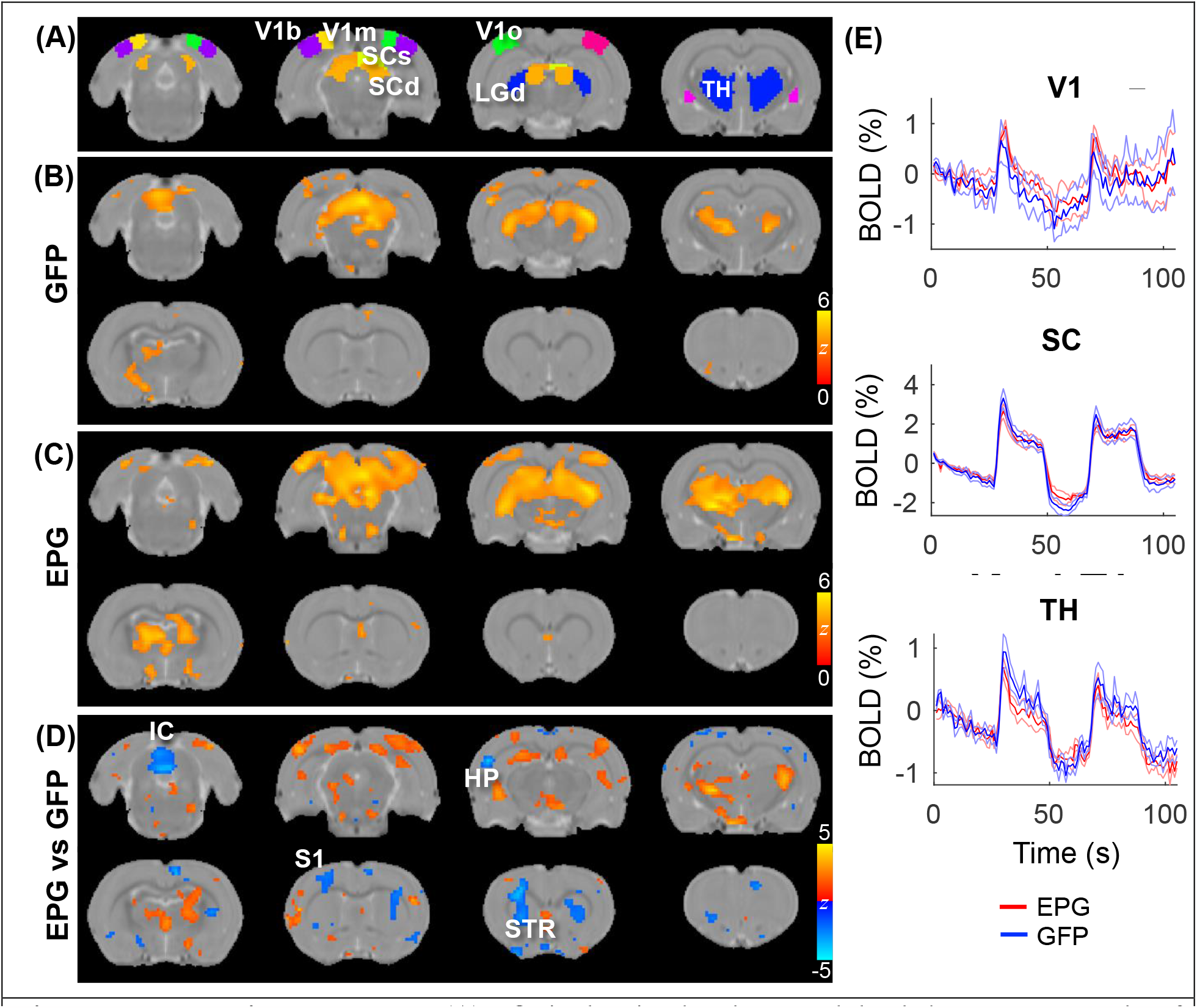
Evoked visual responses. (A) ROI in the visual pathway and the thalamus. Two epochs of 20s visual stimulation were delivered, leading to activation in the visual pathway in the (B) GFP and (C) EPG groups (one-sample t-test, p < 0.01 FDR corrected). (D) The EPG group showed significant increased activation in the primary visual cortex (V1), hippocampus (HP) and ventral thalamus, and decreased activation in the striatum (STR), inferior colliculus (IC) and cerebellum, compared to control rats (two-sample t-test, p<0.05, FDR corrected). (E) Averaged signal time-courses from selected ROI. Error bar represents SEM and the gray bars indicate the stimulation periods.

To further evaluate the effects of visual EPG on other sensory pathways, we conducted electrical forepaw stimulation. As expected, activation in the ventral posterior lateral (VPL) thalamus, and primary somatosensory forelimb area (S1FL) were detected in the GFP-control group (Fig. 4A; p < 0.01 FDR corrected). Interestingly, whereas the activation in the S1FL was comparable, much broader activation in the thalamus, anterior cingulate cortex was seen in the EPG group (Fig. 4B). Comparison between the EPG and control groups revealed increased activation in the TH but decreased activation in the STR, similar to that under visual stimulation.

**Figure 4.**
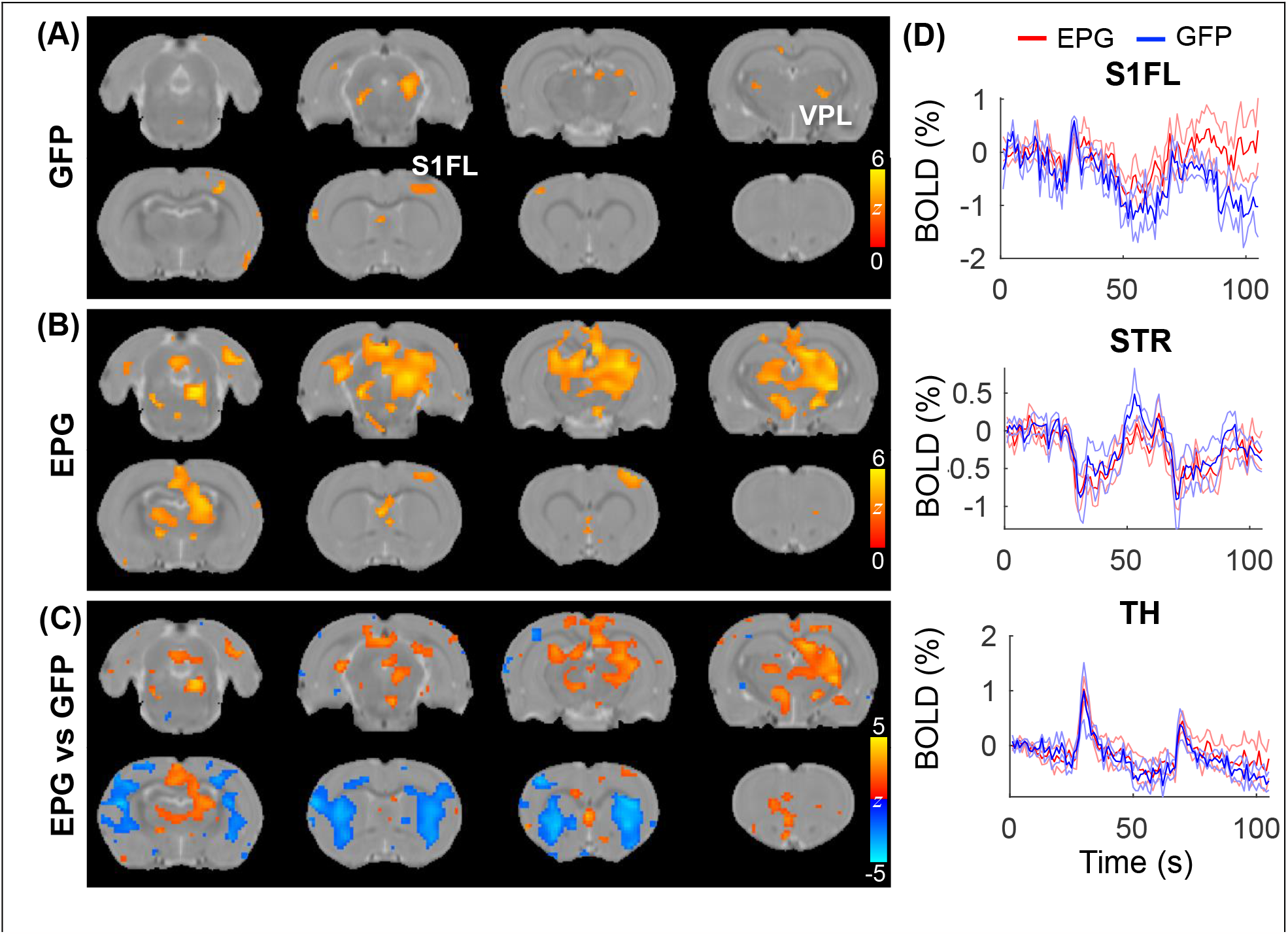
Evoked tactile responses. Two epochs of 20s tactile stimulation were delivered to the forepaw of the rat, leading to significant activation in the somatosensory pathway in the (A) GFP and (B) EPG groups (one-sample t-test, p < 0.01 FDR corrected). (C) Between group comparison shows that the EPG group had increased activation in the Thalamus (TH) and in the anterior cingulate cortex, and decreased activation in the striatum (STR), compared to control rats (two-sample t-test, p<0.05, FDR corrected). (D) Averaged signal time-courses from selected ROI. Errorbar represents SEM and the gray bars indicate the stimulation periods.

Finally, at the end of the fMRI experiments rats were sacrificed for immunohistology validation of EPG expression. Figure 5 shows the expression of EPG (green fluorescence) in excitatory neurons (red fluorescence) in the V1m, indicating wide distribution across the cortical layers.

**Figure 5.**
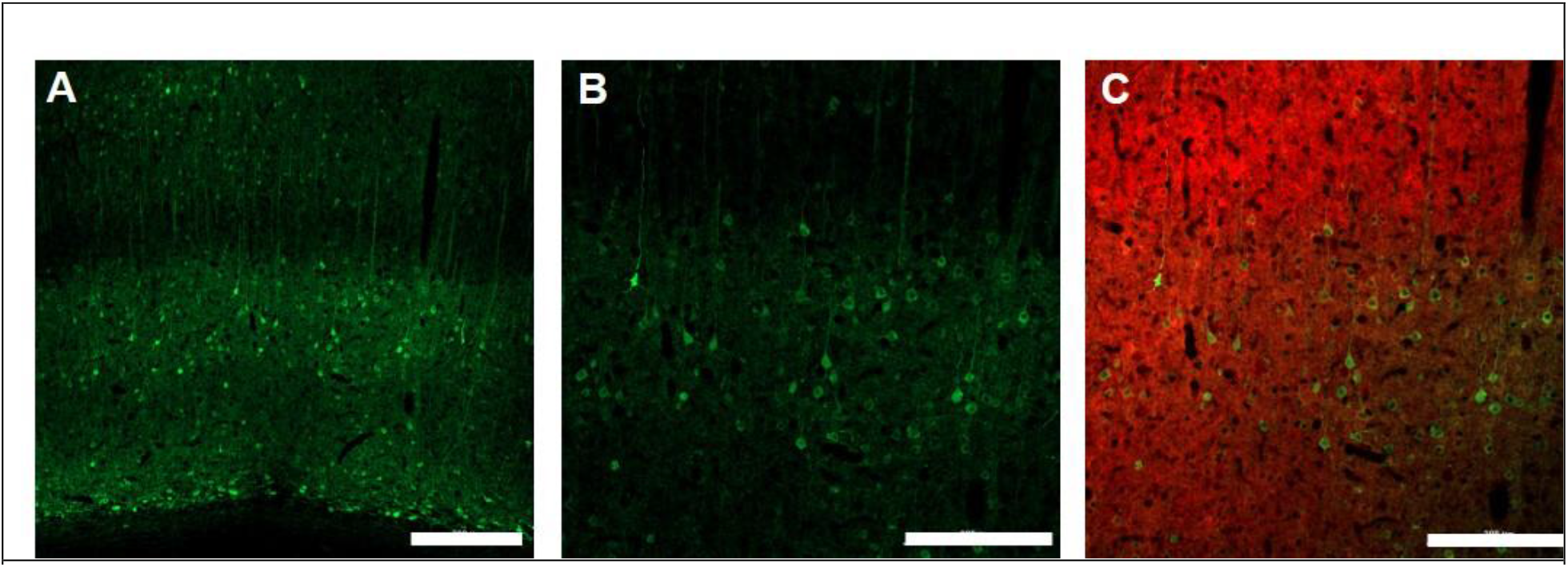
Immunohistology of EPG expression in the visual cortex. 10X magnification ((**A)**, Scale bar = 300µm) and 20X magnification ((**B)**, Scale bar = 200µm of EPG (labeled with green, anti-GFP) throughout the layers of the visual cortex in excitatory neurons (labeled with red, anti-CamKII) ((**C)**, Scale bar=200µm.

## Discussion

Previous work has shown that electromagnetic devices positioned over EPG-expressing neurons induce changes in neural activity *in vivo* and *in vitro*. Here, we demonstrated that EPG can also be activated by strong magnetic fields induced by an MRI, including both the static and the oscillating gradient fields. FC analysis demonstrated that EPG-expressing rats showed extensively stronger FC in sensory areas, and fALFF analysis demonstrated higher signal fluctuation, specifically in the visual areas. These results suggest that by using EPG-based magnetogenetics, MRI can induce changes in the connectivity and activity of neural circuits in rodents. This new approach would allow studying the activity of specific circuits associated with different brain function and disease conditions in a way that was not possible previously.

The molecular structure of the EPG and how it responds to magnetic fields is an active area of research. Strong evidence suggests that exposure to magnetic fields leads to conformational changes in the EPG protein. By fusing split fragments of a certain protein to both termini of the EPG, the fragments can be reassembled into a functional protein under magnetic stimulation due to a conformational change [17]. Electrophysiology recordings in acute brain slices obtained from rats that expressed EPG in the somatosensory cortex show that the time course of neural responses induced by EPG activity is in the order of milliseconds; Magnetic stimulation significantly increased action potentials within 100 ms, and the number of action potentials returned to baseline afterward [19]. Thus, the molecular dynamics of the EPG allow a rapid and controlled activation of neural activity. Indeed, the results from the current fMRI support this hypothesis.

It is essential to determine the precise magnetic field strength required to activate EPG. Studies have consistently demonstrated that exposure to magnetic fields in the mT range can activate a variety of cells and neurons via EPG [14, 17, 20, 21, 27], Nevertheless, no test has been carried out with a stronger magnetic field to date. Indeed, most active proteins are in unstable thermodynamic states and, therefore may not require much energy for the activation [28]. For example, bacterial enzyme kinetics are affected by low magnetic fields via spin-orbit coupling when bound to Xenon gas [29]. Moreover, small magnetic fields (∼3 mT) can affect the enzymatic reaction of horseradish peroxidase via a similar mechanism [30]. These reports indicate that magnetic fields can directly affect proteins by mechanisms other than the movement or alignment of a magnetic core. Furthermore, there is mounting evidence that biochemical reactions can significantly amplify the magnetic field effect [31] and that chemical inhibitors can reverse such effect [32]. Although MRI appears to have a magnetic field strong enough to activate EPG and alter neuronal activity, additional research is required to determine the extent to which the permanent magnetic field of the MRI and the lower, fluctuating magnetic field generated by the EPI gradients contribute to this effect.

In the current study, we started collecting resting-state fMRI because it is possible that just exposing the EPG-expressing rodent to the strong magnetic field of the MRI scanner would activate the EPG. After acquiring twelve minutes of resting-state fMRI we collected evoked response fMRI. Changes in spontaneous neural activity, connectivity, and sensory evoked responses were observed in fALFF, FC, and fMRI analysis. The activity and connectivity at many remote regions from the EPG-expressing site was observed. Based on the axonal connectivity mapping [33, 34], V1 has extensively direct projections to the ACC, RSC, SC, S1, motor cortex and auditory cortex to support visuosensory and visuomotor integration in visually guided behaviors. Therefore, the observed change is mostly likely due the EPG modulation in these downstream pathways. Consistent with the structural and functional connectivity findings, we found somatosensory activation under a tactile stimulation can be modulated by EPG likely via the visuosensory connections. Additionally, the visual and somatosensory cortices directly project to the dorsal striatum to support the sensorimotor functions and action selection [35, 36]. This is consistent with our observation of altered striatal activation under the visual or tactile stimulation though its functional role remains to be elucidated.

Previously, it was demonstrated that EPG can be expressed in different cell types by using specific neural promoters. For example, in a rat model of peripheral nerve injury, EPG was expressed in excitatory neurons in the primary somatosensory cortex under CamKII promoter [21] and in inhibitory interneurons in the hippocampus under hDlx promoter [19]. Like optogenetics and chemogenetics, the EPG can be delivered to any desired brain structure using molecular biology tools, and it can be expressed under cell-specific promoters. In addition, like optogenetics and electronic devices, the activation of cells expressing EPG is immediate. However, unlike optogenetics, chemogenetics, and electronic devices, activation of EPG is minimally invasive and does not require the administration of drugs or implantation of any device. Currently, it does require the delivery of viral vectors through stereotaxic injections into the brain. Nevertheless, new and upcoming approaches for minimally invasive gene delivery that are being designed for different health disorders are likely to offer new ways to deliver transgenes that can cross the blood-brain barrier and target a specific neural population [37-39] and will provide new approaches to deliver EPG into the brain. Thus, magnetogenetics technology eliminates many complications and side effects often associated with current stimulation techniques.

In summary, we demonstrated for the first time a new method to manipulate neural circuits in rodents via magnetic fields inside the MRI. With resting-state and evoked fMRI, we found EPG can modulate downstream pathways activity and FC extending from the visual cortex. This indicates a crucial need of using whole-brain functional imaging to understand downstream effects of a targeted neuromodulation. Further development of the magnetogenetics may provide a new tool for studying connectivity and neural activity in animal models.

## Acknowledgments

the authors would like to acknowledge financial support from the NIH/NINDS: R01-NS098231 (GP and AAG); R01-NS104306 (AAG and GP); UF1NS115817 (GP); NIH/NIBIB: R01-EB031008 (AAG); R01-EB030565 (AAG); R01-EB031936 (AAG and GP); NSF 2027113 (AAG).

## Author contributions

Conceptualization: G.P and A.A.G; Methodology: G.P and C.Q; Analysis; K-H.C and G.P; Writing-Original Draft: G.P, K-H.C, A.A.G; Writing-Review and Editing: K-H.C. G.P, A.A.G, C.Q.

## Data availability

The data that support the findings of this study are available from the corresponding author upon reasonable request.

