## Supporting information for "Magnetogenetic stimulation inside MRI induces spontaneous and evoked changes in neural circuits activity in rats"

**pAAV-CaMKIIα::(EPG(Rat)X3Flag)-IRES-EGFP**

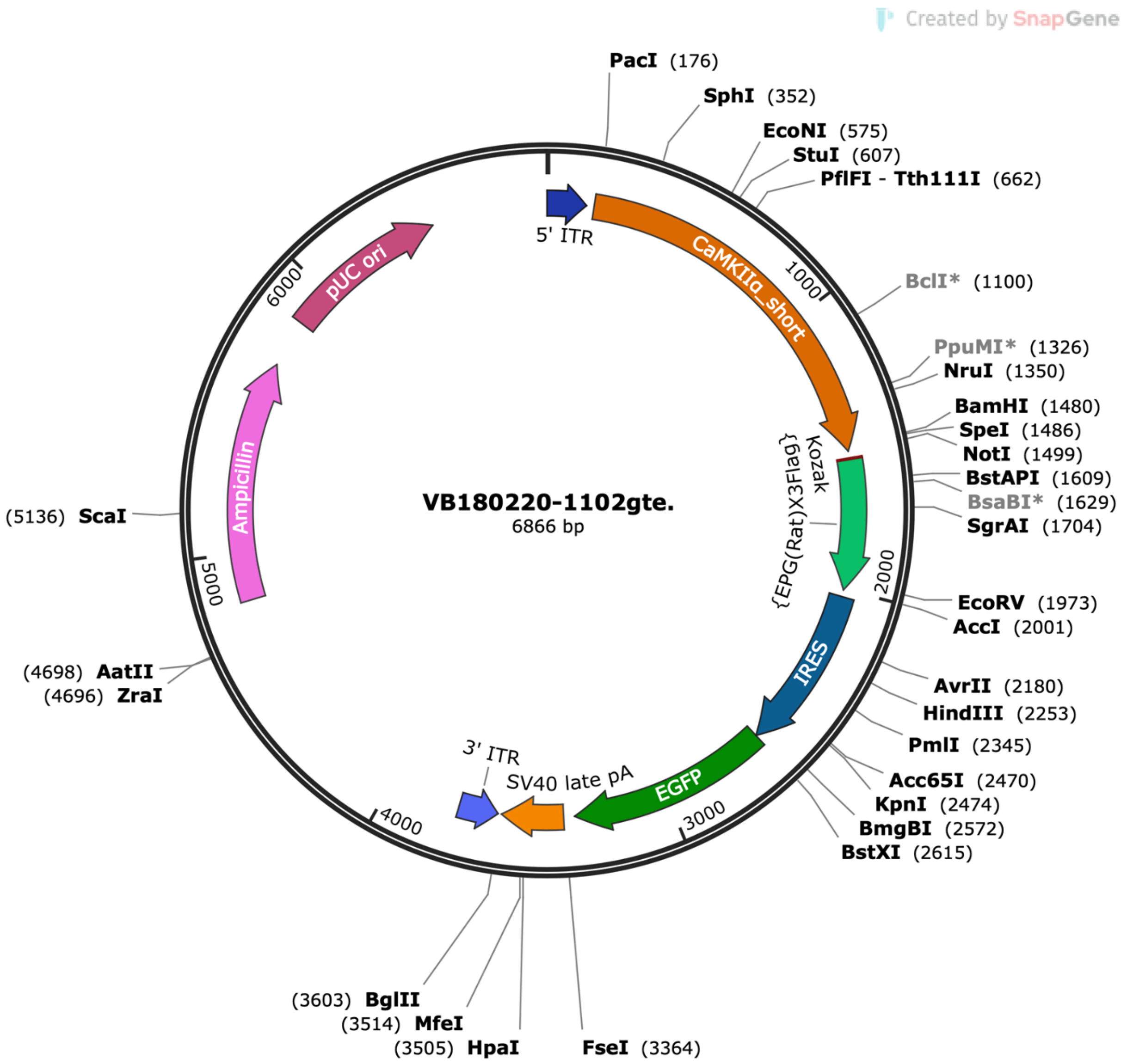


CCTGCAGGCAGCTGCGCGCTCGCTCGCTCACTGAGGCCGCCCGGGCAAAGCCCGGGCGTCGGGCGACCT
TTGGTCGCCCGGCCTCAGTGAGCGAGCGAGCGCGCAGAGAGGGAGTGGCCAACTCCATCACTAGGGGTT
CCTATCGATCAACTTTGTATAGAAAAGTTGCCCTTAATTAACATTATGGCCTTAGGTCACTTCATCTCCATGG
GGTTCTTCTTCTGATTTTCTAGAAAATGAGATGGGGGTGCAGAGAGCTTCCTCAGTGACCTGCCCAGGGT
CACATCAGAAATGTCAGAGCTAGAACTTGAACTCAGATTACTAATCTTAAATTCCATGCCTTGGGGGCATGC
AAGTACGATATACAGAAGGAGTGAACTCATTAGGGCAGATGACCAATGAGTTTAGGAAAGAAGAGTCCAG
GGCAGGGTACATCTACACCACCCGCCCAGCCCTGGGTGAGTCCAGCCACGTTCACCTCATTATAGTTGCCT
CTCTCCAGTCCTACCTTGACGGGAAGCACAAGCAGAAACTGGGACAGGAGCCCCAGGAGACCAAATCTT
CATGGTCCCTCTGGGAGGATGGGTGGGGAGAGCTGTGGCAGAGGCCTCAGGAGGGGCCCTGCTGCTCA
GTGGTGACAGATAGGGGTGAGAAAGCAGACAGAGTCATTCCGTCAGCATTCTGGGTCTGTTTGGTACTTC
TTCTCACGCTAAGGTGGCGGTGTGATATGCACAATGGCTAAAAAGCAGGGAGAGCTGGAAAGAAACAAG

GACAGAGACAGAGGCCAAGTCAACCAGACCAATTCCCAGAGGAAGCAAAGAAACCATTACAGAGACTAC
AAGGGGGAAGGGAAGGAGAGATGAATTAGCTTCCCCTGTAAACCTTAGAACCCAGCTGTTGCCAGGGCA
ACGGGGCAATACCTGTCTCTTCAGAGGAGATGAAGTTGCCAGGGTAACTACATCCTGTCTTTCTCAAGGAC
CATCCCAGAATGTGGCACCCACTAGCCGTTACCATAGCAACTGCCTCTTTGCCCCACTTAATCCCATCCCGTC
TGTTAAAAGGGCCCTATAGTTGGAGGTGGGGGAGGTAGGAAGAGCGATGATCACTTGTGGACTAAGTTT
GTTCGCATCCCCTTCTCCAACCCCCTCAGTACATCACCCTGGGGGAACAGGGTCCACTTGCTCCTGGGCCC
ACACAGTCCTGCAGTATTGTGTATATAAGGCCAGGGCAAAGAGGAGCAGGTTTTAAAGTGAAAGGCAGG
CAGGTGTTGGGGAGGCAGTTACCGGGGCAACGGGAACAGGGCGTTTCGGAGGTGGTTGCCATGGGGAC
CTGGATGCTGACGAAGGCTCGCGAGGCTGTGAGCAGCCACAGTGCCCTGCTCAGAAGCCCCAAGCTCGT
CAGTCAAGCCGGTTCTCCGTTTGCACTCAGGAGCACGGGCAGGCGAGTGGCCCCTAGTTCTGGGGGCAG
CTCTAGAGCGGGGGATCCACTAGTTCTAGAGCGGCCGCCAAGTTTGTACAAAAAAGCAGGCTGCCACCAT
GAAGTGCGTGCTGCTGGGCTTCGCCGCCGTGATCGGCTTCTTCGCCATCGCCGAGAGCCTGACCTGCAAC
ACCTGCAGCGTGAGCCTGATCGGCATCTGCCTGAACCCCGCCACCGCCACCTGCAGCACCAACACCAGCG
TGTGCACCACCGGCAGGGCCAGCTTCACCGGCGTGCTGGGCTTCCTGGGCTTCAACAGCCAGGGCTGCA
CCGAGGGCGCCCAGTGCAACGGCACCGTGAGCGGCAGCATCCTGGGCGCCAGCTACACCGTGACCCAGA
CCTGCTGCAGCACCAACAACTGCAACCCCGTGACCAGCGGCGCCAGCTACGTGCAGATCAGCGTGAGCG
CCGCCCTGAGCGCCGCCCTGCTGGCCTGCGTGTGGGGCCAGAGCGTGTACGACTACAAAGACCATGACG
GTGATTATAAAGATCATGATATCGATTACAAGGATGACGATGACAAGTAGACCCAGCTTTCTTGTACAAAGT
GGGCCCCTCTCCCTCCCCCCCCCCTAACGTTACTGGCCGAAGCCGCTTGGAATAAGGCCGGTGTGCGTTTG
TCTATATGTTATTTTCCACCATATTGCCGTCTTTTGGCAATGTGAGGGCCCGGAAACCTGGCCCTGTCTTCTT
GACGAGCATTCCTAGGGGTCTTTCCCCTCTCGCCAAAGGAATGCAAGGTCTGTTGAATGTCGTGAAGGAA
GCAGTTCCTCTGGAAGCTTCTTGAAGACAAACAACGTCTGTAGCGACCCTTTGCAGGCAGCGGAACCCCC
CACCTGGCGACAGGTGCCTCTGCGGCCAAAAGCCACGTGTATAAGATACACCTGCAAAGGCGGCACAACC
CCAGTGCCACGTTGTGAGTTGGATAGTTGTGGAAAGAGTCAAATGGCTCTCCTCAAGCGTATTCAACAAG
GGGCTGAAGGATGCCCAGAAGGTACCCCATTGTATGGGATCTGATCTGGGGCCTCGGTGCACATGCTTTAC
ATGTGTTTAGTCGAGGTTAAAAAAACGTCTAGGCCCCCCGAACCACGGGGACGTGGTTTTCCTTTGAAAA
ACACGATGATAATATGGCCACAACCATGGTGAGCAAGGGCGAGGAGCTGTTCACCGGGGTGGTGCCCATC
CTGGTCGAGCTGGACGGCGACGTAAACGGCCACAAGTTCAGCGTGTCCGGCGAGGGCGAGGGCGATGC
CACCTACGGCAAGCTGACCCTGAAGTTCATCTGCACCACCGGCAAGCTGCCCGTGCCCTGGCCCACCCTC
GTGACCACCCTGACCTACGGCGTGCAGTGCTTCAGCCGCTACCCCGACCACATGAAGCAGCACGACTTCT
TCAAGTCCGCCATGCCCGAAGGCTACGTCCAGGAGCGCACCATCTTCTTCAAGGACGACGGCAACTACAA
GACCCGCGCCGAGGTGAAGTTCGAGGGCGACACCCTGGTGAACCGCATCGAGCTGAAGGGCATCGACTT
CAAGGAGGACGGCAACATCCTGGGGCACAAGCTGGAGTACAACTACAACAGCCACAACGTCTATATCATG
GCCGACAAGCAGAAGAACGGCATCAAGGTGAACTTCAAGATCCGCCACAACATCGAGGACGGCAGCGTG
CAGCTCGCCGACCACTACCAGCAGAACACCCCCATCGGCGACGGCCCCGTGCTGCTGCCCGACAACCACT
ACCTGAGCACCCAGTCCGCCCTGAGCAAAGACCCCAACGAGAAGCGCGATCACATGGTCCTGCTGGAGTT
CGTGACCGCCGCCGGGATCACTCTCGGCATGGACGAGCTGTACAAGTAACAACTTTATTATACATAGTTGAT
GGCCGGCCGCTTCGAGCAGACATGATAAGATACATTGATGAGTTTGGACAAACCACAACTAGAATGCAGT
GAAAAAAATGCTTTATTTGTGAAATTTGTGATGCTATTGCTTTATTTGTAACCATTATAAGCTGCAATAAACA
AGTTAACAACAACAATTGCATTCATTTTATGTTTCAGGTTCAGGGGGAGGTGTGGGAGGTTTTTTAAAGCA
AGTAAAACCTCTACAAATGTGGTAATCGATAGATCTAGGAACCCCTAGTGATGGAGTTGGCCACTCCCTCTC
TGCGCGCTCGCTCGCTCACTGAGGCCGGGCGACCAAAGGTCGCCCGACGCCCGGGCTTTGCCCGGGCGG
CCTCAGTGAGCGAGCGAGCGCGCAGCTGCCTGCAGGCAGCTTGGCACTGGCCGTCGTTTTACAACGTCG
TGACTGGGAAAACCCTGGCGTTACCCAACTTAATCGCCTTGCAGCACATCCCCCTTTCGCCAGCTGGCGTA

ATAGCGAAGAGGCCCGCACCGATCGCCCTTCCCAACAGTTGCGCAGCCTGAATGGCGAATGGCGCCTGAT
GCGGTATTTTCTCCTTACGCATCTGTGCGGTATTTCACACCGCATACGTCAAAGCAACCATAGTACGCGCCC
TGTAGCGGCGCATTAAGCGCGGCGGGTGTGGTGGTTACGCGCAGCGTGACCGCTACACTTGCCAGCGCC
CTAGCGCCCGCTCCTTTCGCTTTCTTCCCTTCCTTTCTCGCCACGTTCGCCGGCTTTCCCCGTCAAGCTCTAA
ATCGGGGGCTCCCTTTAGGGTTCCGATTTAGTGCTTTACGGCACCTCGACCCCAAAAAACTTGATTTGGGT
GATGGTTCACGTAGTGGGCCATCGCCCTGATAGACGGTTTTTCGCCCTTTGACGTTGGAGTCCACGTTCTT
TAATAGTGGACTCTTGTTCCAAACTGGAACAACACTCAACCCTATCTCGGGCTATTCTTTTGATTTATAAGGG
ATTTTGCCGATTTCGGCCTATTGGTTAAAAAATGAGCTGATTTAACAAAAATTTAACGCGAATTTTAACAAA
ATATTAACGTTTACAATTTTATGGTGCACTCTCAGTACAATCTGCTCTGATGCCGCATAGTTAAGCCAGCCCC
GACACCCGCCAACACCCGCTGACGCGCCCTGACGGGCTTGTCTGCTCCCGGCATCCGCTTACAGACAAGC
TGTGACCGTCTCCGGGAGCTGCATGTGTCAGAGGTTTTCACCGTCATCACCGAAACGCGCGAGACGAAAG
GGCCTCGTGATACGCCTATTTTTATAGGTTAATGTCATGATAATAATGGTTTCTTAGACGTCAGGTGGCACTT
TTCGGGGAAATGTGCGCGGAACCCCTATTTGTTTATTTTTCTAAATACATTCAAATATGTATCCGCTCATGAG
ACAATAACCCTGATAAATGCTTCAATAATATTGAAAAAGGAAGAGTATGAGTATTCAACATTTCCGTGTCGC
CCTTATTCCCTTTTTTGCGGCATTTTGCCTTCCTGTTTTTGCTCACCCAGAAACGCTGGTGAAAGTAAAAGA
TGCTGAAGATCAGTTGGGTGCACGAGTGGGTTACATCGAACTGGATCTCAACAGCGGTAAGATCCTTGAG
AGTTTTCGCCCCGAAGAACGTTTTCCAATGATGAGCACTTTTAAAGTTCTGCTATGTGGCGCGGTATTATCC
CGTATTGACGCCGGGCAAGAGCAACTCGGTCGCCGCATACACTATTCTCAGAATGACTTGGTTGAGTACTC
ACCAGTCACAGAAAAGCATCTTACGGATGGCATGACAGTAAGAGAATTATGCAGTGCTGCCATAACCATGA
GTGATAACACTGCGGCCAACTTACTTCTGACAACGATCGGAGGACCGAAGGAGCTAACCGCTTTTTTGCA
CAACATGGGGGATCATGTAACTCGCCTTGATCGTTGGGAACCGGAGCTGAATGAAGCCATACCAAACGAC
GAGCGTGACACCACGATGCCTGTAGCAATGGCAACAACGTTGCGCAAACTATTAACTGGCGAACTACTTAC
TCTAGCTTCCCGGCAACAATTAATAGACTGGATGGAGGCGGATAAAGTTGCAGGACCACTTCTGCGCTCG
GCCCTTCCGGCTGGCTGGTTTATTGCTGATAAATCTGGAGCCGGTGAGCGTGGGTCTCGCGGTATCATTGC
AGCACTGGGGCCAGATGGTAAGCCCTCCCGTATCGTAGTTATCTACACGACGGGGAGTCAGGCAACTATG
GATGAACGAAATAGACAGATCGCTGAGATAGGTGCCTCACTGATTAAGCATTGGTAACTGTCAGACCAAGT
TTACTCATATATACTTTAGATTGATTTAAAACTTCATTTTTAATTTAAAAGGATCTAGGTGAAGATCCTTTTTG
ATAATCTCATGACCAAAATCCCTTAACGTGAGTTTTCGTTCCACTGAGCGTCAGACCCCGTAGAAAAGATCA
AAGGATCTTCTTGAGATCCTTTTTTTCTGCGCGTAATCTGCTGCTTGCAAACAAAAAAACCACCGCTACCAG
CGGTGGTTTGTTTGCCGGATCAAGAGCTACCAACTCTTTTTCCGAAGGTAACTGGCTTCAGCAGAGCGCA
GATACCAAATACTGTTCTTCTAGTGTAGCCGTAGTTAGGCCACCACTTCAAGAACTCTGTAGCACCGCCTAC
ATACCTCGCTCTGCTAATCCTGTTACCAGTGGCTGCTGCCAGTGGCGATAAGTCGTGTCTTACCGGGTTGGA
CTCAAGACGATAGTTACCGGATAAGGCGCAGCGGTCGGGCTGAACGGGGGGTTCGTGCACACAGCCCAG
CTTGGAGCGAACGACCTACACCGAACTGAGATACCTACAGCGTGAGCTATGAGAAAGCGCCACGCTTCCC
GAAGGGAGAAAGGCGGACAGGTATCCGGTAAGCGGCAGGGTCGGAACAGGAGAGCGCACGAGGGAG
CTTCCAGGGGGAAACGCCTGGTATCTTTATAGTCCTGTCGGGTTTCGCCACCTCTGACTTGAGCGTCGATTT
TTGTGATGCTCGTCAGGGGGGCGGAGCCTATGGAAAAACGCCAGCAACGCGGCCTTTTTACGGTTCCTGG
CCTTTTGCTGGCCTTTTGCTCACATGTTCTTTCCTGCGTTATCCCCTGATTCTGTGGATAACCGTATTACCGCC
TTTGAGTGAGCTGATACCGCTCGCCGCAGCCGAACGACCGAGCGCAGCGAGTCAGTGAGCGAGGAAGC
GGAAGAGCGCCCAATACGCAAACCGCCTCTCCCCGCGCGTTGGCCGATTCATTAATGCAGCTGGCACGAC
AGGTTTCCCGACTGGAAAGCGGGCAGTGAGCGCAACGCAATTAATGTGAGTTAGCTCACTCATTAGGCAC
CCCAGGCTTTACACTTTATGCTTCCGGCTCGTATGTTGTGTGGAATTGTGAGCGGATAACAATTTCACACAG
GAAACAGCTATGACCATGATTACGAATTG

### pAAV-CaMKIIα::EGFP

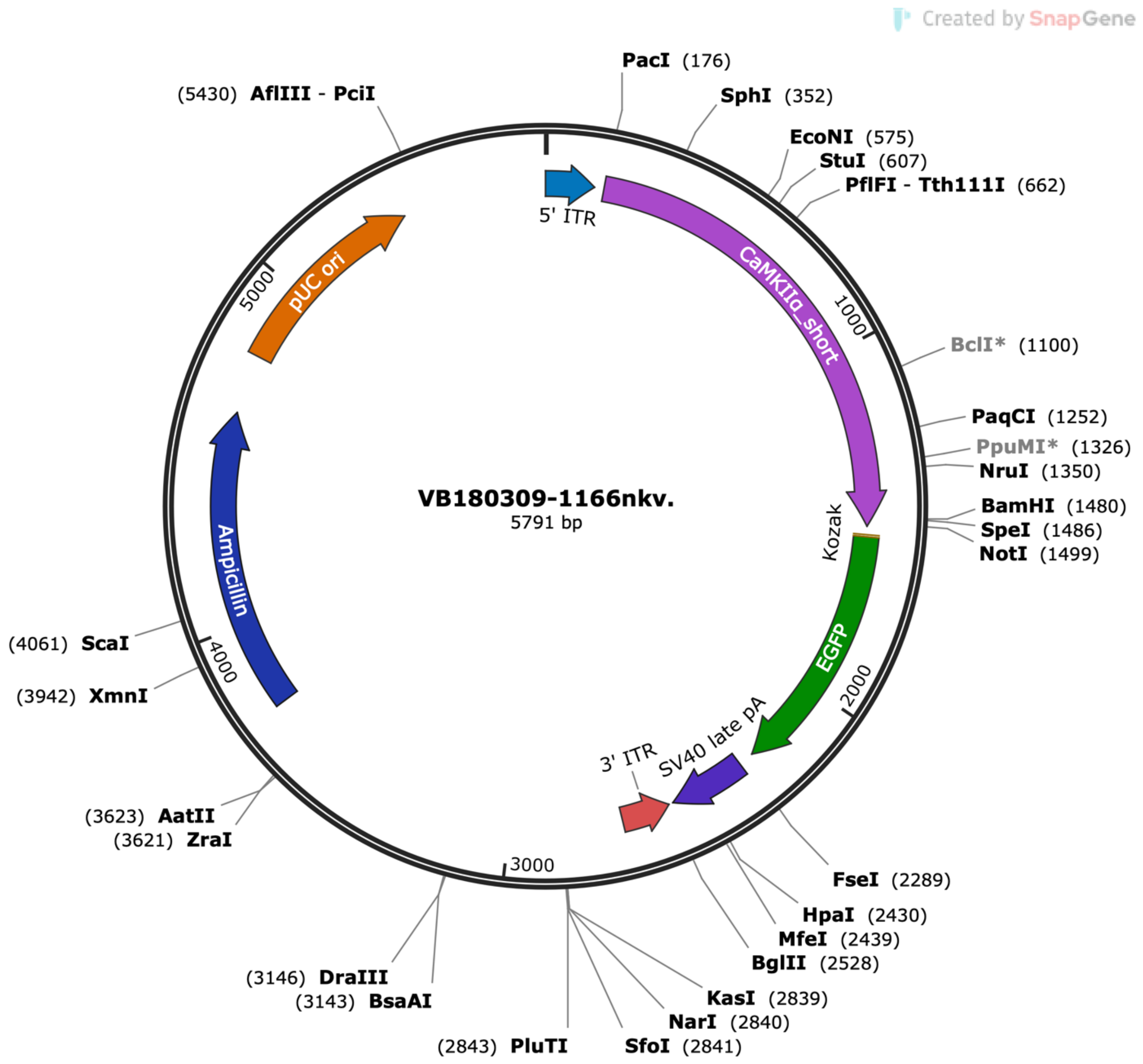


CCTGCAGGCAGCTGCGCGCTCGCTCGCTCACTGAGGCCGCCCGGGCAAAGCCCGGGCGTCGGGCGACCT
TTGGTCGCCCGGCCTCAGTGAGCGAGCGAGCGCGCAGAGAGGGAGTGGCCAACTCCATCACTAGGGGTT
CCTATCGATCAACTTTGTATAGAAAAGTTGCCCTTAATTAACATTATGGCCTTAGGTCACTTCATCTCCATGG
GGTTCTTCTTCTGATTTTCTAGAAAATGAGATGGGGGTGCAGAGAGCTTCCTCAGTGACCTGCCCAGGGT
CACATCAGAAATGTCAGAGCTAGAACTTGAACTCAGATTACTAATCTTAAATTCCATGCCTTGGGGGCATGC
AAGTACGATATACAGAAGGAGTGAACTCATTAGGGCAGATGACCAATGAGTTTAGGAAAGAAGAGTCCAG
GGCAGGGTACATCTACACCACCCGCCCAGCCCTGGGTGAGTCCAGCCACGTTCACCTCATTATAGTTGCCT
CTCTCCAGTCCTACCTTGACGGGAAGCACAAGCAGAAACTGGGACAGGAGCCCCAGGAGACCAAATCTT
CATGGTCCCTCTGGGAGGATGGGTGGGGAGAGCTGTGGCAGAGGCCTCAGGAGGGGCCCTGCTGCTCA
GTGGTGACAGATAGGGGTGAGAAAGCAGACAGAGTCATTCCGTCAGCATTCTGGGTCTGTTTGGTACTTC
TTCTCACGCTAAGGTGGCGGTGTGATATGCACAATGGCTAAAAAGCAGGGAGAGCTGGAAAGAAACAAG
GACAGAGACAGAGGCCAAGTCAACCAGACCAATTCCCAGAGGAAGCAAAGAAACCATTACAGAGACTAC

AAGGGGGAAGGGAAGGAGAGATGAATTAGCTTCCCCTGTAAACCTTAGAACCCAGCTGTTGCCAGGGCA
ACGGGGCAATACCTGTCTCTTCAGAGGAGATGAAGTTGCCAGGGTAACTACATCCTGTCTTTCTCAAGGAC
CATCCCAGAATGTGGCACCCACTAGCCGTTACCATAGCAACTGCCTCTTTGCCCCACTTAATCCCATCCCGTC
TGTTAAAAGGGCCCTATAGTTGGAGGTGGGGGAGGTAGGAAGAGCGATGATCACTTGTGGACTAAGTTT
GTTCGCATCCCCTTCTCCAACCCCCTCAGTACATCACCCTGGGGGAACAGGGTCCACTTGCTCCTGGGCCC
ACACAGTCCTGCAGTATTGTGTATATAAGGCCAGGGCAAAGAGGAGCAGGTTTTAAAGTGAAAGGCAGG
CAGGTGTTGGGGAGGCAGTTACCGGGGCAACGGGAACAGGGCGTTTCGGAGGTGGTTGCCATGGGGAC
CTGGATGCTGACGAAGGCTCGCGAGGCTGTGAGCAGCCACAGTGCCCTGCTCAGAAGCCCCAAGCTCGT
CAGTCAAGCCGGTTCTCCGTTTGCACTCAGGAGCACGGGCAGGCGAGTGGCCCCTAGTTCTGGGGGCAG
CTCTAGAGCGGGGGATCCACTAGTTCTAGAGCGGCCGCCAAGTTTGTACAAAAAAGCAGGCTGCCACCAT
GGTGAGCAAGGGCGAGGAGCTGTTCACCGGGGTGGTGCCCATCCTGGTCGAGCTGGACGGCGACGTAA
ACGGCCACAAGTTCAGCGTGTCCGGCGAGGGCGAGGGCGATGCCACCTACGGCAAGCTGACCCTGAAGT
TCATCTGCACCACCGGCAAGCTGCCCGTGCCCTGGCCCACCCTCGTGACCACCCTGACCTACGGCGTGCAG
TGCTTCAGCCGCTACCCCGACCACATGAAGCAGCACGACTTCTTCAAGTCCGCCATGCCCGAAGGCTACGT
CCAGGAGCGCACCATCTTCTTCAAGGACGACGGCAACTACAAGACCCGCGCCGAGGTGAAGTTCGAGGG
CGACACCCTGGTGAACCGCATCGAGCTGAAGGGCATCGACTTCAAGGAGGACGGCAACATCCTGGGGCA
CAAGCTGGAGTACAACTACAACAGCCACAACGTCTATATCATGGCCGACAAGCAGAAGAACGGCATCAAG
GTGAACTTCAAGATCCGCCACAACATCGAGGACGGCAGCGTGCAGCTCGCCGACCACTACCAGCAGAAC
ACCCCCATCGGCGACGGCCCCGTGCTGCTGCCCGACAACCACTACCTGAGCACCCAGTCCGCCCTGAGCA
AAGACCCCAACGAGAAGCGCGATCACATGGTCCTGCTGGAGTTCGTGACCGCCGCCGGGATCACTCTCGG
CATGGACGAGCTGTACAAGTAAACCCAGCTTTCTTGTACAAAGTGGTGATGGCCGGCCGCTTCGAGCAGA
CATGATAAGATACATTGATGAGTTTGGACAAACCACAACTAGAATGCAGTGAAAAAAATGCTTTATTTGTGA
AATTTGTGATGCTATTGCTTTATTTGTAACCATTATAAGCTGCAATAAACAAGTTAACAACAACAATTGCATTC
ATTTTATGTTTCAGGTTCAGGGGGAGGTGTGGGAGGTTTTTTAAAGCAAGTAAAACCTCTACAAATGTGGT
AATCGATAGATCTAGGAACCCCTAGTGATGGAGTTGGCCACTCCCTCTCTGCGCGCTCGCTCGCTCACTGA
GGCCGGGCGACCAAAGGTCGCCCGACGCCCGGGCTTTGCCCGGGCGGCCTCAGTGAGCGAGCGAGCGC
GCAGCTGCCTGCAGGCAGCTTGGCACTGGCCGTCGTTTTACAACGTCGTGACTGGGAAAACCCTGGCGTT
ACCCAACTTAATCGCCTTGCAGCACATCCCCCTTTCGCCAGCTGGCGTAATAGCGAAGAGGCCCGCACCGA
TCGCCCTTCCCAACAGTTGCGCAGCCTGAATGGCGAATGGCGCCTGATGCGGTATTTTCTCCTTACGCATCT
GTGCGGTATTTCACACCGCATACGTCAAAGCAACCATAGTACGCGCCCTGTAGCGGCGCATTAAGCGCGGC
GGGTGTGGTGGTTACGCGCAGCGTGACCGCTACACTTGCCAGCGCCCTAGCGCCCGCTCCTTTCGCTTTCT
TCCCTTCCTTTCTCGCCACGTTCGCCGGCTTTCCCCGTCAAGCTCTAAATCGGGGGCTCCCTTTAGGGTTCC
GATTTAGTGCTTTACGGCACCTCGACCCCAAAAAACTTGATTTGGGTGATGGTTCACGTAGTGGGCCATCG
CCCTGATAGACGGTTTTTCGCCCTTTGACGTTGGAGTCCACGTTCTTTAATAGTGGACTCTTGTTCCAAACT
GGAACAACACTCAACCCTATCTCGGGCTATTCTTTTGATTTATAAGGGATTTTGCCGATTTCGGCCTATTGGT
TAAAAAATGAGCTGATTTAACAAAAATTTAACGCGAATTTTAACAAAATATTAACGTTTACAATTTTATGGTG
CACTCTCAGTACAATCTGCTCTGATGCCGCATAGTTAAGCCAGCCCCGACACCCGCCAACACCCGCTGACG
CGCCCTGACGGGCTTGTCTGCTCCCGGCATCCGCTTACAGACAAGCTGTGACCGTCTCCGGGAGCTGCAT
GTGTCAGAGGTTTTCACCGTCATCACCGAAACGCGCGAGACGAAAGGGCCTCGTGATACGCCTATTTTTAT
AGGTTAATGTCATGATAATAATGGTTTCTTAGACGTCAGGTGGCACTTTTCGGGGAAATGTGCGCGGAACC
CCTATTTGTTTATTTTTCTAAATACATTCAAATATGTATCCGCTCATGAGACAATAACCCTGATAAATGCTTCAA
TAATATTGAAAAAGGAAGAGTATGAGTATTCAACATTTCCGTGTCGCCCTTATTCCCTTTTTTGCGGCATTTT
GCCTTCCTGTTTTTGCTCACCCAGAAACGCTGGTGAAAGTAAAAGATGCTGAAGATCAGTTGGGTGCACG
AGTGGGTTACATCGAACTGGATCTCAACAGCGGTAAGATCCTTGAGAGTTTTCGCCCCGAAGAACGTTTTC

CAATGATGAGCACTTTTAAAGTTCTGCTATGTGGCGCGGTATTATCCCGTATTGACGCCGGGCAAGAGCAA
CTCGGTCGCCGCATACACTATTCTCAGAATGACTTGGTTGAGTACTCACCAGTCACAGAAAAGCATCTTACG
GATGGCATGACAGTAAGAGAATTATGCAGTGCTGCCATAACCATGAGTGATAACACTGCGGCCAACTTACT
TCTGACAACGATCGGAGGACCGAAGGAGCTAACCGCTTTTTTGCACAACATGGGGGATCATGTAACTCGC
CTTGATCGTTGGGAACCGGAGCTGAATGAAGCCATACCAAACGACGAGCGTGACACCACGATGCCTGTAG
CAATGGCAACAACGTTGCGCAAACTATTAACTGGCGAACTACTTACTCTAGCTTCCCGGCAACAATTAATAG
ACTGGATGGAGGCGGATAAAGTTGCAGGACCACTTCTGCGCTCGGCCCTTCCGGCTGGCTGGTTTATTGC
TGATAAATCTGGAGCCGGTGAGCGTGGGTCTCGCGGTATCATTGCAGCACTGGGGCCAGATGGTAAGCCC
TCCCGTATCGTAGTTATCTACACGACGGGGAGTCAGGCAACTATGGATGAACGAAATAGACAGATCGCTGA
GATAGGTGCCTCACTGATTAAGCATTGGTAACTGTCAGACCAAGTTTACTCATATATACTTTAGATTGATTTA
AAACTTCATTTTTAATTTAAAAGGATCTAGGTGAAGATCCTTTTTGATAATCTCATGACCAAAATCCCTTAAC
GTGAGTTTTCGTTCCACTGAGCGTCAGACCCCGTAGAAAAGATCAAAGGATCTTCTTGAGATCCTTTTTTT
CTGCGCGTAATCTGCTGCTTGCAAACAAAAAAACCACCGCTACCAGCGGTGGTTTGTTTGCCGGATCAAG
AGCTACCAACTCTTTTTCCGAAGGTAACTGGCTTCAGCAGAGCGCAGATACCAAATACTGTTCTTCTAGTGT
AGCCGTAGTTAGGCCACCACTTCAAGAACTCTGTAGCACCGCCTACATACCTCGCTCTGCTAATCCTGTTAC
CAGTGGCTGCTGCCAGTGGCGATAAGTCGTGTCTTACCGGGTTGGACTCAAGACGATAGTTACCGGATAA
GGCGCAGCGGTCGGGCTGAACGGGGGGTTCGTGCACACAGCCCAGCTTGGAGCGAACGACCTACACCG
AACTGAGATACCTACAGCGTGAGCTATGAGAAAGCGCCACGCTTCCCGAAGGGAGAAAGGCGGACAGGT
ATCCGGTAAGCGGCAGGGTCGGAACAGGAGAGCGCACGAGGGAGCTTCCAGGGGGAAACGCCTGGTAT
CTTTATAGTCCTGTCGGGTTTCGCCACCTCTGACTTGAGCGTCGATTTTTGTGATGCTCGTCAGGGGGGCG
GAGCCTATGGAAAAACGCCAGCAACGCGGCCTTTTTACGGTTCCTGGCCTTTTGCTGGCCTTTTGCTCACA
TGTTCTTTCCTGCGTTATCCCCTGATTCTGTGGATAACCGTATTACCGCCTTTGAGTGAGCTGATACCGCTCG
CCGCAGCCGAACGACCGAGCGCAGCGAGTCAGTGAGCGAGGAAGCGGAAGAGCGCCCAATACGCAAAC
CGCCTCTCCCCGCGCGTTGGCCGATTCATTAATGCAGCTGGCACGACAGGTTTCCCGACTGGAAAGCGGG
CAGTGAGCGCAACGCAATTAATGTGAGTTAGCTCACTCATTAGGCACCCCAGGCTTTACACTTTATGCTTCC
GGCTCGTATGTTGTGTGGAATTGTGAGCGGATAACAATTTCACACAGGAAACAGCTATGACCATGATTACG
AATTG

**Supplementary Table S1**
The full list of gray matter regions in each hemisphere used in the ROI analysis.

| ROI | Abbreviation |
| --- | --- |
| Agranular Dysgranular Insular Cortex | ADI |
| Agranular Insular Cortex | AI |
| Amygdalohyppocampic Area | AHi |
| Amygdalopiriform Cortex | APir |
| Basal Forebrain Region | BF |
| Bed Nucleus of the Stria Terminalis | BNST |
| Cornu Ammonis 1 | CA1 |
| Cornu Ammonis 2 | CA2 |
| Cornu Ammonis 3 | CA3 |
| Dentate Gyrus | DG |
| Dorso Lateral Orbital Cortex | LO |
| Dysgranular Insular Cortex | DI |
| Ectorhinal Cortex | Ect |
| Entorhinal Cortex | Ent |
| Fasciola Cinereum | FaC |
| Frontal Association Cortex | FrA |
| Globus Pallidus | GP |
| Glomerular Layer of the Accessory Olfactory Bulb | AOB |
| Glomerular Layer of the Olfactory Bulb | OB |
| Lateral Temporal Associative Cortex | TeAL |
| Lateral Primary Auditory Cortex | Au1 |
| Granule Cell Level of the Cerebellum | CeG |
| Hypothalamic Region | Hy |
| Interpeduncular Nucleus | IPN |
| Lateral Entorhinal Cortex Internal part | LEntIn |
| Lateral Entorhinal Cortex | LEnt |
| Lateral Entorhinal Cortex external part | LEntEx |
| Lateral Parietal Associative Cortex | LPtA |
| Lateral Secondary Visual Cortex | V2L |
| Medial Entorhinal Cortex | Ment |
| Medial Parietal Associative Cortex | MPtA |
| Medio Lateral Secondary Visual Cortex | V2ML |
| Medio Medial Secondary Visual Cortex | V2MM |
| Molecular Cell Level of the Cerebellum | CeM |
| Olfactory Bulb | OB |
| Orbitofrontal Region | OFC |
| Parasubiculum | PaS |
| Parietal Cortex Postero Caudal Part | PtPC |
| Parietal Cortex Postero Dorsal Part | PtPD |
| Parietal Cortex Postero Rostral | PtPR |

| | |
|---|---|
| Periaqueductal Gray | PAG |
| Perirhinal Area 35 | PRh35 |
| Perirhinal Area 36 | PRh36 |
| Perirhinal Cortex | PRh |
| Posterior Agralunar Insular Cortex | AIP |
| PreLimbic System | PrL |
| Presubiculum | PrS |
| Pretectal Region | PrT |
| Primary Auditory Cortex | AUD |
| Primary Cingular Cortex | Cg1 |
| Primary Motor Cortex | M1 |
| Primary Somatosensory Cortex Barrel field | S1BF |
| Primary Somatosensory Cortex Dysgranular | S1DZ |
| Primary Somatosensory Cortex Dysgranular Zone 0 | S1DZ0 |
| Primary Somatosensory Cortex Forelimb | S1FL |
| Primary Somatosensory Cortex Hindlimb | S1HL |
| Primary Somatosensory Cortex Jaw | S1J |
| Primary Somatosensory Cortex | S1C |
| Primary Somatosensory Cortex Shoulder | S1Sh |
| Primary Somatosensory Cortex Trunk | S1Tr |
| Primary Somatosensory Cortex Upperlips | S1ULp |
| Primary Visual Cortex Binocular Area | V1b |
| Primary Visual Cortex | V1c |
| Primary Visual Cortex Monocular Area | V1m |
| Retosplenial Dysgranular Cortex | RSCd |
| Retrosplenial Granular Cortex Part A | RSCa |
| Retrosplenial Granular Cortex Part B | RSCb |
| Secondary Auditory Cortex Dorsal Part | Au2D |
| Secondary Auditory Cortex Ventral Part | Au2V |
| Secondary Cingular Cortex | Cg2 |
| Secondary Motor Cortex | M2 |
| Secondary Somatosensory Cortex | S2 |
| Striatum | STR |
| Subiculum | Sub |
| Substantia Nigra | SN |
| Superficial Gray Layer of the Superior Colliculus | SCs |
| Temporal Associative Cortex | TeA |
| Brainstem | BS |
| Deeper Layers of the Superior Colliculus | SCd |
| External Cortex of the Inferior Colliculus | ICe |
| Septal Region | Sep |
| Subthalamic Nucleus | STN |
| Thalamus | TH |

| | |
|---|---|
| Periventricular Grey | PVG |
| Pons | Pons |
